# Spectrally and temporally resolved estimation of neural signal diversity

**DOI:** 10.1101/2023.03.30.534922

**Authors:** Pedro A.M. Mediano, Fernando E. Rosas, Andrea I. Luppi, Valdas Noreika, Anil K. Seth, Robin L. Carhart-Harris, Lionel Barnett, Daniel Bor

## Abstract

Quantifying the complexity of neural activity has provided fundamental insights into cognition, consciousness, and clinical conditions. However, the most widely used approach to estimate the complexity of neural dynamics, Lempel-Ziv complexity (LZ), has fundamental limitations that substantially restrict its domain of applicability. In this article we leverage the information-theoretic foundations of LZ to overcome these limitations by introducing a complexity estimator based on state-space models —which we dub *Complexity via State-space Entropy Rate* (CSER). While having a performance equivalent to LZ in discriminating states of consciousness, CSER boasts two crucial advantages: 1) CSER offers a principled decomposition into spectral components, which allows us to rigorously investigate the relationship between complexity and spectral power; and 2) CSER provides a temporal resolution two orders of magnitude better than LZ, which allows complexity analyses of e.g. event-locked neural signals. As a proof of principle, we use MEG, EEG and ECoG datasets of humans and monkeys to show that CSER identifies the gamma band as the main driver of complexity changes across states of consciousness; and reveals early entropy increases that *precede* the standard ERP in an auditory mismatch negativity paradigm by approximately 20ms. Overall, by overcoming the main limitations of LZ and substantially extending its range of applicability, CSER opens the door to novel investigations on the fine-grained spectral and temporal structure of the signal complexity associated with cognitive processes and conscious states.

## Introduction

### Complexity measures in neuroscience

Since the advent of complexity science, there has been great interest in its application to understand the brain (***Turkheimer et al., 2022***). Lempel and Ziv’s now-classical complexity measure (***Lempel and Ziv, 1976***) has been highly influential and widely applied in the neuroimaging literature, especially for the neuroscientific study of consciousness. LZ complexity is a simple scalar metric that has been consistently shown to be a robust indicator of depth of anaesthesia (***Zhang et al., 2001***), disorders of consciousness (***Casali et al., 2013***), sleep (***Abásolo et al., 2015***; ***Schartner et al., 2017b***), and, more recently, the psychedelic state (***Schartner et al., 2017a***; ***Timmermann et al., 2019***; ***Mediano et al., 2020b***), making it one of the most effective known functional markers of conscious state in humans. LZ is also effective as a biomarker of mental disorders such as schizophrenia (***Ibáñez-Molina et al., 2018***; ***Rajpal et al., 2022***) and depression (***Bachmann et al., 2015***; ***Akar et al., 2015***), as well as tracking fluctuations of normal wakefulness related to drowsiness (***Mediano et al., 2021***), mind wandering (***Ibáñez-Molina and Iglesias-Parro, 2014***), or artistic improvisation (***Dolan et al., 2018***).

Despite its practical effectiveness and widespread use, LZ has longstanding limitations that severely restrict its domain of applicability. First, the application of LZ requires that the data be discretised, which loses information and introduces artificial non-linearities (***Ibáñez-Molina et al., 2015***; ***Yeh and Shi, 2018***). Second, the LZ algorithm needs to be provided with relatively large windows of data (on the order of a few seconds for typical EEG data), which limits its temporal resolution —making it impossible to do complexity analyses of non-stationary data, such as the fast time-locked neural events studied with ERP signals. Third, it is not possible to do a principled spectral decomposition of LZ, a fundamental obstacle to our understanding of the relationship between complexity and power spectrum —which is highly relevant in neuroscience as electrophysiological signals are often differently distributed across several frequency bands.

In this paper we introduce a novel approach to calculate an LZ-style complexity that overcomes all these limitations. To do so, our approach builds on the rich literature of *state-space models*, powerful and versatile statistical time series models widely used in neuroscience and beyond (***Durbin and Koopman, 2012***). Our method, which we call *Complexity via State-space Entropy Rate* (CSER), solves all of LZ’s above-mentioned issues:

- CSER does not require the signal to be discretized, allowing it to fully exploit continuous signals and avoiding potential artefacts introduced by the discretization procedure.
- CSER allows instantaneous (i.e. sample-by-sample) estimation, bringing complexity analyses to the realm of event-related paradigms.
- CSER has a principled spectral decomposition, closing the gap between complexity and traditional spectral analyses.

In the rest of the paper, we first describe the basic intuitions behind the common use of LZ in neuroscience. We then introduce our estimator, CSER, and discuss its tight mathematical links with signal diversity and prediction error (***Den Ouden et al., 2012***). Next, we showcase CSER’s properties in a comparison of different states of consciousness and in data from an auditory mismatch negativity paradigm. Finally, we conclude with a discussion on how to interpret complexity measures applied to neural activity, and note on the applicability of CSER to other neuroimaging modalities.

### Complexity and entropy rate

Before we delve into the formulation of our new estimator and the new results obtained with it, it is worth examining the theoretical foundations of LZ as a measure of complexity.

The informational content of a discrete signal 𝒳 = (*X*_1_, *X*_2_, …) at time *t* can be quantified by its entropy,

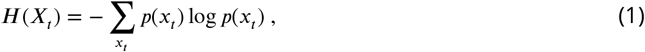

which can be seen as measuring the difficulty of guessing the signal’s value at that point in time—the more difficult this is, the more information one gains by learning the signal’s actual value. If the signal preferentially takes some specific values, then the signal is easier to predict, which corresponds to a low entropy. On the other hand, if the signal takes all of its available values with similar frequency (i.e. its histogram is nearly flat), then there is high uncertainty about its current value and hence observing it will be very informative, which corresponds to a case of high entropy.

Crucially, entropy only takes into account the relative frequency of values, but not the order in which they appear. As an example, consider a signal A corresponding to the sequence 01010101, and a signal B, corresponding to the sequence 01110010. A naive plug-in estimator (***Panzeri et al., 2007***) applied to these signals would yield the same entropy for both, as they have the same frequency of 0s and 1s. However, if one knows the past trajectory of the sequence it is clearly much easier to predict the next value of signal A than signal B. This aspect of a signal’s unpredictability —the difficulty of predicting the next value *after knowing all previous ones* —is quantified by the signal’s *entropy rate*, here denoted by *h*(𝒳). In the above example, signals A and B have the same entropy, but signal A has lower entropy rate than B. In other words, entropy is invariant to reshuffling the order of elements in a sequence, whereas entropy rate is not —making the latter a more appropriate measure of dynamical complexity.

Estimating entropy rate in practice from limited data is, however, a challenging task. This is due to several factors, most prominently the *“*curse of dimensionality*”* (***Paninski, 2003***). This is where the LZ algorithm comes into play.

In essence, the original algorithm by ***Lempel and Ziv*** (***1976***) breaks up a signal into patterns, and uses the number of distinct patterns to quantify the complexity of that signal. Regular sequences with repeating patterns have low LZ, and rich signals with many patterns have high LZ. A fundamental result by ***Ziv*** (***1978***) shows that this number of patterns can be used to efficiently estimate the entropy rate of the process that generated the data. Specifically, given a discrete stochastic process^1^ 𝒳= (*X*_1_, *X*_2_, …, *X*_*T*_) with LZ complexity *c*(𝒳), its entropy rate can be estimated as^2^

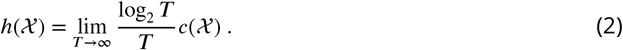

Furthermore, the concept of entropy rate allows us to draw a direct connection between the common interpretation of LZ as signal diversity and another important framework in neuroscience: *predictive processing*. In essence, the more diverse a signal is, the higher the prediction error one would incur when trying to forecast its future (see Methods for details).

In this work, we provide evidence that the empirical successes of LZ can be linked directly to the underlying and more general mathematical construct of entropy rate. This opens a way to formulate metrics, based on entropy rate, that are similar to LZ in spirit but do not suffer from the same limitations —as we showcase in the remainder of this article.

## Results

Having outlined the core principles behind LZ, and also having described its inherent limitations, we can now introduce our novel estimator, CSER. We demonstrate that CSER preserves the utility of LZ while offering multiple clear advantages over it, illustrated across three sets of results. First, we show that CSER preserves the valuable empirical effectiveness of LZ, and can clearly discriminate between states of consciousness, in different species and different neuroimaging modalities. Then, we illustrate two examples of new analysis methods that can be performed by leveraging the advantageous properties of state-space models, which are not possible with LZ: spectral and temporal decomposition of entropy in neural signals. Mathematical details of CSER and validation analyses on artificial data can be found in the Methods section.

### Estimating entropy rate via state-space models

The core principle of state-space analysis is to assume that the observed data, 𝒳, can be modelled as noisy observations of a hidden process 𝒵 that is not accessible to direct measurement (Fig. 1). Mathematically, a state-space model is fully specified by two ingredients: the dynamics of the hidden state 𝒵 (horizontal arrows); and the observation process that relates 𝒵 with the observed data 𝒳 (vertical arrows). A simple yet effective choice is to assume that both the dynamics and observation processes are linear and normally distributed. Fitting a state-space model, then, corresponds to finding the linear coefficients and covariance matrices that best describe the relationship between *z*_*t*_ and *z*_*t*+1_, and between *z*_*t*_ and *x*_*t*_ (see Methods for more details).

The properties of state-space models make it straightforward to calculate entropy rate from a fitted model. In particular, the key property of state-space models is that the hidden state *z*_*t*_ con-tains all the relevant information about *x*_1_ …, *x*_*t*−1_ needed to predict *x*_*t*_. Then, thanks to the assumption of normality and the link between entropy rate and prediction error, entropy rate can be calculated with the usual formula of the entropy of a Gaussian distribution (***Cover and Thomas, 2006***, Sec. 8.4) —resulting in the CSER estimator. See the Methods section for details on state-space models, model selection, and the robustness of CSER to model misspecification.

**Figure 1.**
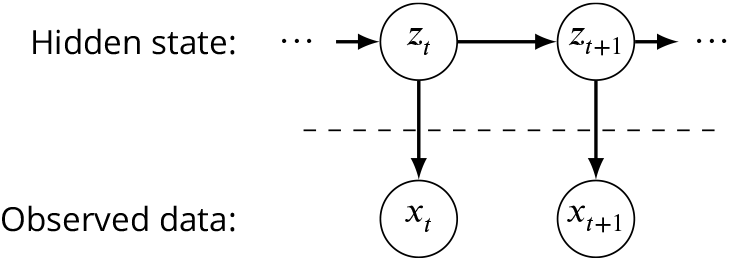
Diagram of a state-space process.

### Discriminative power between states of consciousness

The main feature that makes LZ complexity interesting for neuroscientists is its empirical predictive value: for instance, it consistently decreases in states of loss of consciousness such as sleep or general anaesthesia (***Zhang et al., 1999***; ***Casali et al., 2013***; ***Varley et al., 2020***);^3^ and increases in states with richer subjective contents of consciousness like the psychedelic state (***Schartner et al., 2017a***; ***Mediano et al., 2020b***). Therefore, one would expect that any candidate competing measure of complexity should preserve those desirable empirical properties.

To comprehensively validate this hypothesis, we analyse data of subjects in different well-defined states of consciousness obtained through different neuroimaging modalities. In particular, we use three datasets:

1. MEG data from 15 human subjects under the effects of the psychedelic drug LSD, as well as a placebo (PLA) (***Carhart-Harris et al., 2016***).
2. EEG data from 9 human subjects in dreamless NREM sleep, as well as a wakeful rest baseline (***Wong et al., 2020***).
3. ECoG data from 4 macaque subjects under the effects of ketamine and medetomidine (KTMD) anaesthesia, as well as a wakeful rest baseline (***Yanagawa et al., 2013***).

In all cases, we compute CSER and LZ (for comparison) for each channel separately following the procedure outlined in the Methods section, and report results averaged across all channels (Fig. 2). Results show that CSER fully agrees with LZ, increasing in subjects under the effects of LSD, and decreasing in subjects in NREM sleep or general anaesthesia. The effects have the same sign for LZ and CSER across all three datasets, and are significant with *p <* 0.001 for the psychedelics and sleep datasets, and with *p <* 0.05 for the anaesthesia dataset (details in Supplementary Table 1). Furthermore, the spatial distributions across the brain for LZ and CSER show qualitative agreement, especially for the modalities with good spatial resolution (MEG and ECoG). These results confirm that CSER is able to perform the same discriminatory role as LZ on a wide range of datasets and states. We next highlight two clear advantages of using CSER instead of LZ: spectral decomposition and temporal resolution.

**Figure 2.**
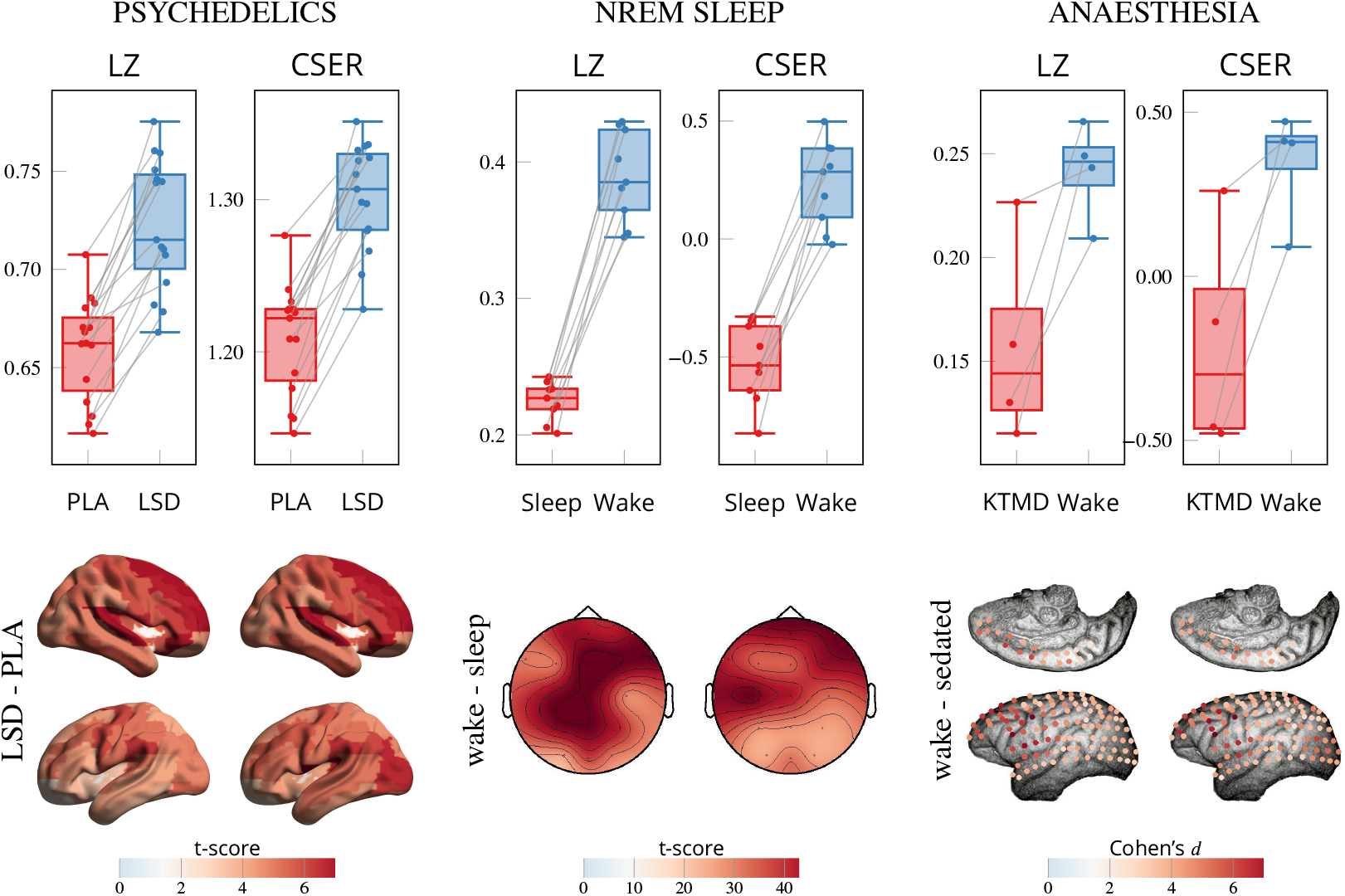
CSER discriminates between states of consciousness. That is, it preserves the key empirical properties of LZ: entropy increases under the effects of psychedelics (*left*), and decreases in NREM sleep (*middle*) and general anaesthesia (*right*). Top row shows subject-level averages, bottom row shows spatial distributions of LZ and CSER. Given the different locations of ECoG sensors in each subject of the anaesthesia dataset, we show only one subject (Chibi) and use Cohen’s *d* instead of *t*-scores.

### Spectral decomposition of entropy changes

Given the increasing prominence of LZ analyses in neuroscience, multiple efforts have been made to elucidate whether, and to what extent, changes in LZ across states of consciousness relate to neural activity in different frequency bands (***Schartner et al., 2017b***; ***Boncompte et al., 2021***). Answering this question, however, requires sophisticated surrogate data tests (***Mediano et al., 2020a***), and definite answers are yet to be established.

In contrast, the relation between linear models and spectral decomposition is well known (***Hannan and Deistler, 2012***), which allows us to perform an exact, analytical decomposition of the entropy rate into frequency components (see Methods for details). Therefore, one can split the complexity of a signal into different frequency bands, with the guarantee that the terms associated with bands that cover the whole spectrum will sum up to the broadband CSER. This allows us to properly interpret their relative and absolute magnitudes, and to attribute changes in entropy rate between conditions to a particular band in a principled manner.^4^

As a proof of concept, we applied this spectral decomposition to the three datasets used in the previous section using the standard partition of the spectrum into frequency bands (Fig. 3). Given the guarantee that differences in all spectral components sum up to the total difference, we can conclude that the changes in complexity across states of consciousness are mainly driven (in absolute magnitude) by high-frequency neural activity.

Our results provide formal validation of the long-standing hypotheses linking high-frequency and non-rhythmic signals to consciousness. Gamma oscillations were at the center of early consciousness work by ***Crick and Koch*** (***1990***),^5^ and have been posited as the neural underpinning of self-awareness (***Lou et al., 2017***). They also align with more recent evidence showing the power of gamma activity in discriminating between states of consciousness (***Walter and Hinterberger, 2022***), as well as with results by ***Canales-Johnson et al. (2019***) showing that it is broadband (rather than rhythmic) components that encode prediction errors, which in turn have been put forward as a tool to understand conscious contents (***Hohwy and Seth, 2020***). However, we emphasise that our results are distinct from the known results of decreased gamma coherence with loss of consciousness (e.g. ***Cavinato et al., 2015***) —even when controlled for average gamma coherence, differences in CSER between conscious states remain qualitatively unchanged (Supp. Table 3).

Interestingly, CSER’s ability to decompose the complexity of a signal into frequency bands can give us new insights by effectively providing more *“*dimensions*”* to study the complexity of neural data. As an example, note that in the comparison between LSD and placebo (Fig. 3, left), all bands except gamma actually have a small, but significant, negative change in CSER, *opposite* to the overall trend. Future research should disentangle the mechanistic origins of these different complexity changes, and explore if they have different effects on behaviour and subjective experience.

**Figure 3.**
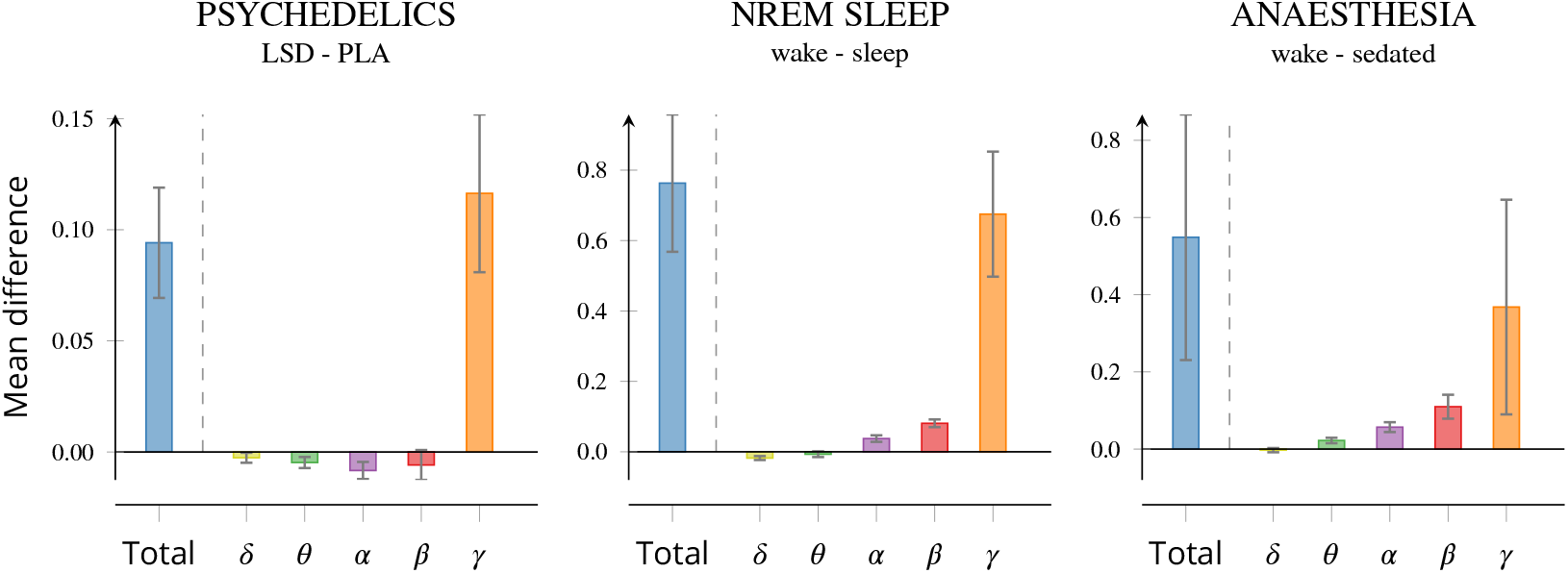
Spectral decomposition of CSER differences across states of consciousness reveals largest contribution from *γ* frequencies. Plots show mean and standard deviation of subject-level differences in CSER decomposed into frequency bands. Note that in each plot the sum of complexity in all bands equals the total (or broadband) complexity. In all three cases, the largest contribution to the difference is high-frequency neural activity. Frequency bands used are *δ*: 1–4 Hz, *θ*: 4–8 Hz, *α*: 8–14 Hz, *β*: 14–25 Hz, *γ*: >25 Hz.

### Temporally-resolved entropy measures

So far, LZ has been considered a measure of *state* —i.e. a property of the typical dynamics observed in the resting-state spontaneous neural activity of a subject. A few recent studies have attempted to capture a more fine-grained temporal evolution of LZ, typically by using a sliding window, although such approaches suffer from a trade-off between time-resolution and estimation noise. Hence, this method only works when the process under consideration has comparatively slow dynamics.^6^

This limitation of seeing LZ only as a *“*state measure*”* is caused, at least in part, by the requirement of large sequence lengths for enabling a robust estimation (a few thousand samples, or several seconds for EEG; c.f. Fig. 5). However, leveraging the explicit parametric form of the state-space models that CSER is based on, we can formulate an instantaneous entropy rate that provides an analogue of LZ computable *for each time point* in multi-trial data.

To illustrate this powerful capability of CSER, we analyse ECoG data from macaque monkeys undergoing an auditory oddball task (***Komatsu et al., 2015***). The auditory oddball is a passive listening task in which monkeys listen to one of two stimuli: a *‘*standard’ tone which is consistent with prior expectation, and a *‘*deviant’ tone which isn’t (Fig. 4, left). One of the most characteristic and widely studied phenomena related to this task is the presence of a *mismatch negativity* (MMN) reponse in the event-related potential (ERP): a strong negative peak following the deviant stimulus (Fig. 4, top right). The MMN occurs approximately 50 to 100 ms post-stimulus in marmosets (***Komatsu et al., 2015***) and 150 to 250 ms in humans (***Näätänen et al., 2007***), and has been studied in a wide variety of experimental settings.

Importantly, one of the leading theories of the mechanisms behind MMN describes it as representing a violation of the brain’s predictions of incoming sensory signals —in essence, when processing the deviant stimulus the brain incurs a *‘*prediction error’ that results in stronger activity (***Garrido et al., 2009***). Given that higher entropy rate is directly linked with a larger prediction error (see Methods), we hypothesised that the deviant stimulus would also elicit an increase in entropy rate (with respect to the standard stimulus) as measured by CSER.

To study the time-resolved entropy rate of this data, we employed the following procedure:

i) Fit a state-space model to the baseline pre-stimulus data, obtaining a set of model parameters.
ii) Use the obtained parameters to evaluate the model and make one-step-ahead predictions in standard and deviant time series, estimating the residual time series.

With the estimated residuals and the known baseline residual variance we can compute a local (i.e. instantaneous) entropy rate, analogous to the model prediction error at time *t* (see Methods and Appendix for details). Using this method, we can compare the instantaneous entropy during the course of standard and deviant percepts in the macaque ECoG, and compare them to the known ERP traces (Fig. 4, right).

In agreement with our predictions, deviant stimuli induce a significant increase in prediction error (and thus, entropy rate) with respect to standard stimuli. Furthermore, and perhaps more interestingly, the peak difference in instantaneous entropy rate *precedes* the ERP peak by approximately 20 ms. We speculate that this highly temporally localised entropy peak could represent the onset of the prediction error itself, that steers neural dynamics in different trajectories depending on the nature of the stimulus, while the difference in neural activity (i.e. ERP amplitude) reflects the spontaneous evolution of these two trajectories.^7^ Naturally, we do not make strong generalisation claims for this phenomenon based on a single subject, although we believe these results warrant further study of the temporal entropy profile of other prediction error-related tasks.

**Figure 4.**
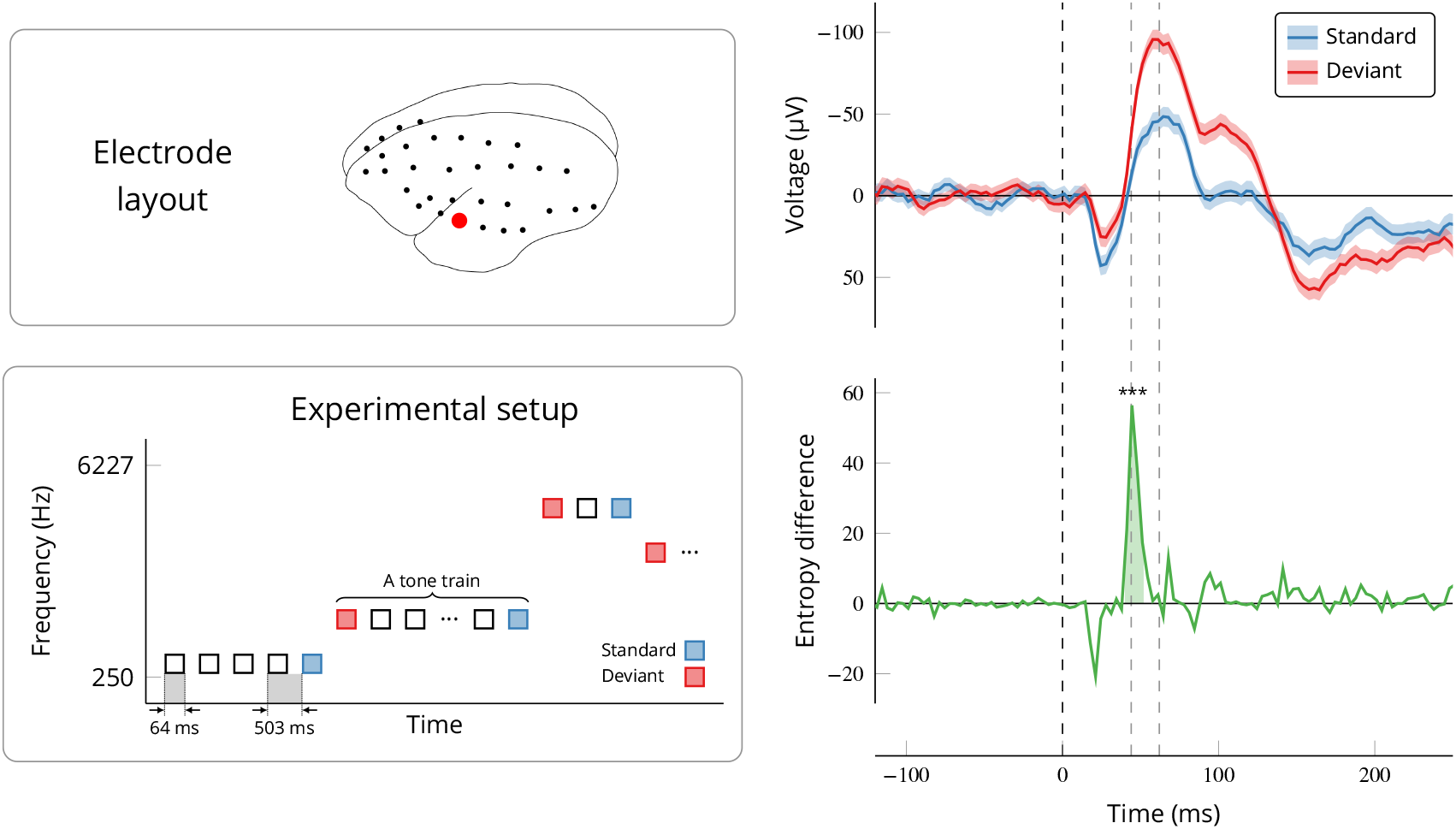
Time-resolved entropy estimates differentiate between time courses of standard and deviant percepts. (*top left*) Layout of ECoG electrodes overlaid on the monkey’s cortex, with the selected electrode in red. (*bottom left*) Schematic diagram of experimental paradigm, in which the subject listens to a tone train composed of *‘*standard’ and *‘*deviant’ tones (see text for details). (*right*) Event-related potentials of standard and deviant trials (*top*), and the instantaneous entropy rate difference computed via CSER (*bottom*). Stars represent a significant cluster with *p <* 0.001. Note that the entropy difference precedes the ERP by approximately 20 ms. Original data from ***Komatsu et al. (2015***) and the [Neurotycho](http://www.neurotycho.org/) database.

**Figure 5.**
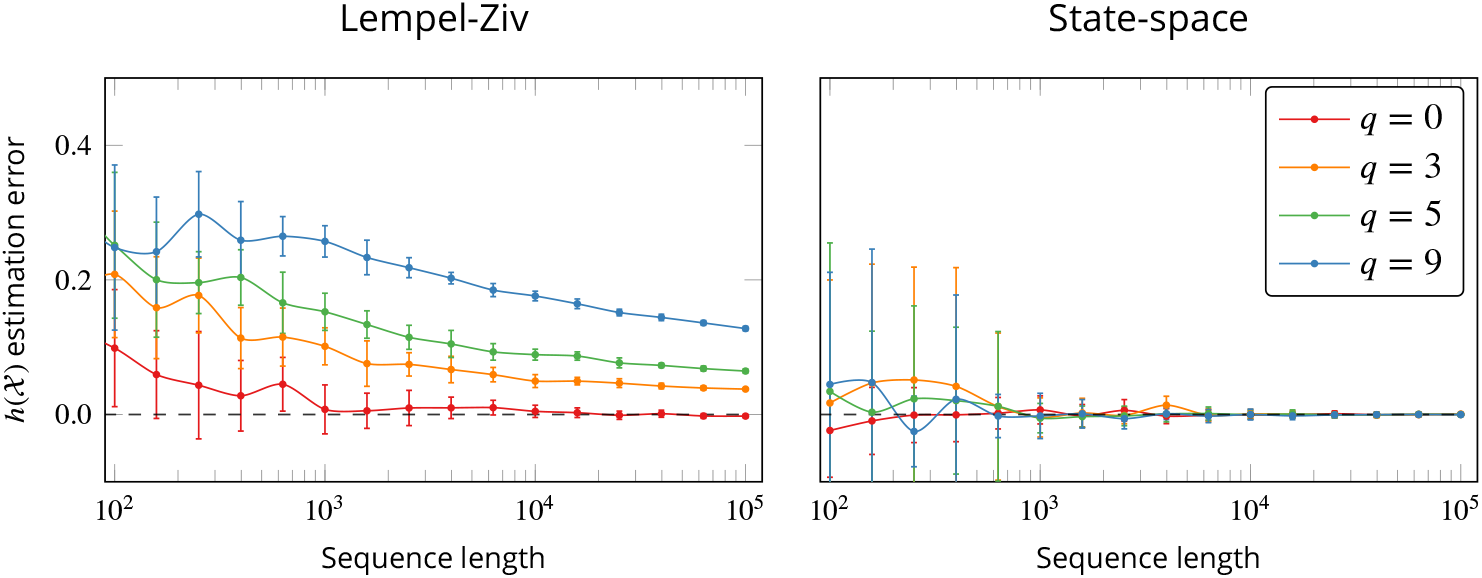
LZ and CSER approximate entropy rate, but CSER converges faster. (*left*) LZ-estimated entropy rates of discrete signals with different lengths (x-axis) and memory order *q*. Except for low values of *q*, LZ shows a slow convergence to the true entropy rate (black line). (*right*) Similar analysis using CSER and synthetic real-valued signals. CSER converges to the true entropy rate for all values of *q* within approximately 1000 samples, within the range of typical M/EEG datasets. Note the logarithmic scale of the x-axis

## Discussion

Electrophysiological data such as EEG, MEG and ECoG, have as their main advantage over other brain-scanning techniques, such as fMRI, that they provide temporally rich information about neuronal activity in different frequency bands. Over the last two decades, the most common method to examine the complexity of these signals has been Lempel-Ziv complexity (LZ), which can be seen as an estimator of the information-theoretic concept of entropy rate. Despite the remarkable practical efficacy of LZ, it is —by construction —unable to give information about the temporal or spectral distribution of this complexity.

The present work introduces a new information-theoretic tool to cognitive neuroscience, CSER, which has distinct advantages over LZ. First, we have shown that CSER has substantially better temporal resolution compared to LZ and is highly sensitive to changes in cognitive state on a mismatched negativity task, potentially detecting a cognitively important neural signal before standard ERPs do. Second, unlike LZ, CSER naturally provides a principled spectral decomposition, yielding intriguing insights about the relationship between gamma-band activity and changes in conscious state.

Across species and imaging modalities, we demonstrate the ability of CSER to provide information about both of these aspects, which LZ could simply not provide. These results may serve as a proof of concept, opening the door to a wide range of new investigations of spectral and temporal aspects of neural complexity observed on multiple kinds of neuroimaging data in cognitive and consciousness studies.

### Advantages and limitations of state-space models in neuroimaging

Throughout this paper we have showcased the power of state-space time series models for a variety of neuroimaging data analyses. Our choice of state-space as models is due to their generality and flexibility, since *i*) they are easy to use and estimate, and do not require discretising the data, *ii*) they are robust to noise and misspecifications in the fitting process, and *iii*) they are able to model and account for a variety of noise factors common in neural data.^8^

Furthermore, the rich analysis possibilities enabled by state-space modelling (as evidenced by the spectral and temporal decompositions above) enable us to explore new dimensions describing the complexity of neural dynamics —which could possibly be mapped to different aspects of consciousness and cognition (***Luppi et al., 2021***). Additionally, the possibility of having sample-by-sample entropy rate estimates allows for the application of multiple analysis techniques from the field of local information dynamics, such as instantaneous information transfer (***Lizier, 2010***) or integrated information (***Mediano et al., 2022***). Conceptually, one could see the generation of entropy rate time series from data as a transformation *“*from volts to bits,*”* opening the door to new analyses on entropy time-series complementary to those on conventional ERPs. We believe these techniques could reveal new information about neural processes and could be a fruitful avenue of future research.

At the same time, however, there are some specific circumstances where state-space models need to be applied with care. Two of these are *i*) applications to fMRI data, and *ii*) M/EEG data after ICA component removal (or average re-referencing). In the case of fMRI, the problem can be related to the haemodynamic response filter, which may introduce singularities in the data (***Solo, 2016***),^9^ and may be alleviated by restricting the maximum model order and regression horizon hyperparameters during the fitting process (see Methods for details). When dealing with preprocessed M/EEG data, ICA component removal may reduce the rank of the data and incur numerical errors when fitting a single state-space model to large sets of channels. In this case, one could mitigate the problem by fitting separate models to smaller subsets of channels, reducing the regression horizon, or adding a very small amount of white noise to the data.

Finally, note that state-space models are not the only alternative to LZ to estimate entropy rate: there exist other parametric methods, such as auto-regressive models (***Barnett and Seth, 2014***), as well as non-parametric methods, such as spectral factorisation (***Chand and Dhamala, 2014***). Nonetheless, state-space models remain our model of choice due to their greater tractability and flexibility, which makes them readily usable in most neuroscientific contexts. It is also possible to build more sophisticated state-space modes, e.g. by combining them with multi-taper techniques (***Kim et al., 2018***).

### Concluding remarks

In this paper we have presented a new method, which we call *Complexity via State-space Entropy Rate* (CSER), as a principled estimator of signal diversity for electrophysiological time series. We have shown that CSER has the desirable empirical properties of successful complexity measures, like Lempel-Ziv complexity, while substantially extending Lempel-Ziv’s capabilities in several ways. Combining four datasets comprising three distinct neuroimaging modalities, several states of consciousness, and an auditory task, we have shown that CSER is a valuable analysis tool that can provide spectrally and temporally resolved insights that were previously impossible —in the first case showing that the difference in complexity across states of consciousness is attributable to high-frequency activity; and in the latter, showing that instantaneous complexity peaks *before* the standard ERP signature of mismatch negativity. To make this method widely available for the neuroscience community, we provide an open-source implementation of CSER and other entropy rate estimators for several programming languages in https://www.github.com/pmediano/EntRate.

Overall, these results emphasise the neuroscientific value of a principled information-theoretic approach, helping us distil the key properties of known methods while empowering neuroscientists to investigate previously inaccessible dimensions of their data.

## Methods

### Datasets and preprocessing

In order to benchmark CSER against LZ, we decided to include data spanning both *i*) multiple neuroimaging modalities, and *ii*) a wide range of states of consciousness. With this in mind, the results in Figs. 2 and 3 were obtained with the following data:

### Psychedelics

We use the MEG data first reported by ***Carhart-Harris et al. (2016***) of *N* = 15 subjects after an infusion of intravenous LSD (or a placebo). Data were source-reconstructed to the centroids of each region in the Automated Anatomical Labelling (AAL) atlas (***Tzourio-Mazoyer et al., 2002***) using a standard LCMV beamformer. For a full description of the source-reconstruction pipeline see ***Mediano et al. (2020b***).

### NREM sleep

We used the EEG data of *N* = 9 subjects during sleep, some of which were previously reported by ***Wong et al. (2020***). Although the original study focused on the neurophysiology of dreams, here we used only segments of data from dreamless NREM sleep, and compared it against a wakeful rest baseline.

### Anaesthesia

We used the ECoG data of *N* = 4 marmoset monkeys sedated with KTMD anaesthesia first reported by ***Yanagawa et al. (2013***). Data were obtained from the open access [Neurotycho](http://www.neurotycho.org/) database and divided into *‘*awake’ and *‘*sedated’ periods.

For all datasets, in addition to modality-specific cleaning, we filtered the data using a lowpass filter with a 100 Hz cutoff, removed line noise by subtracting a least-squares-fit sinusoidal signal at 50 Hz and harmonics, and downsampled it to 200 Hz. For both LZ and CSER, we segmented the data into pseudo-stationary epochs, computed LZ and CSER for each epoch and channel (sources in the LSD dataset, electrodes in the others), and averaged across all epochs and channels. To compute LZ, each epoch was further detrended and binarised around its mean.

Finally, for the time-resolved analysis of an auditory oddball paradigm we use data previously presented by ***Komatsu et al. (2015***). We used only data from the monkey Fr, since it is the only one publicly available in the NeuroTycho website. Given that ECoG data is typically less corrupted by noise, and that for the analysis in Fig. 4 we were interested in the fine temporal structure of the process, we relaxed the lowpass filter to 150 Hz and the downsampling to 300 Hz.

### The two faces of LZ: signal diversity and intrinsic prediction error

Mathematically, the entropy rate of a stochastic process 𝒳 is defined as the asymptotic rate of growth of its entropy, i.e.

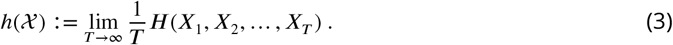

This expression refers to the joint entropy of full trajectories (*X*_1_, …, *X*_*T*_), denoted by *H*(*X*_1_, …, *X*_*T*_). This quantity is related to how much the signal is exploring possible paths: the number of trajectories of *T* time-steps that are effectively visited by the system is approximately 2^*T h*(𝒳)^.^10^ Hence, for sequences built on *K* different symbols, the signal is exploring a fraction 2^*h*(𝒳)^/*K* of the space of possible configurations. These results give the basis for a rigorous interpretation of LZ as *signal diversity*: a higher LZ implies that the system explores a larger fraction of its possible trajectories.

On the other hand, standard results in information theory (***Cover and Thomas, 2006***, Th. 4.2.1) state that the entropy rate can also be expressed as the entropy of the present state conditioned on its past:

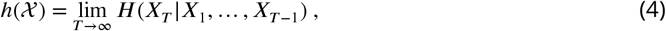

i.e. the uncertainty in the moment-to-moment prediction of the next state of the system. In Gaussian systems this corresponds to the logarithm of the mean squared error (***Cover and Thomas, 2006***), and in discrete systems it is related to the probability of misclassification (***Fano, 1961***; ***Feder and Merhav, 1994***). Thus, entropy rate is also formally linked with *intrinsic prediction error*:^11^ a higher LZ value means it is harder to predict the next value of the signal —even with complete knowledge of its past trajectory.

Therefore, thanks to information theory and the concept of entropy rate we can bridge between two previously disconnected interpretations of complexity in neural dynamics: signal diversity and prediction error. We believe rigorous investigation of the mathematics underlying analysis frameworks can lead to more convergence between neuroscientific theories (***Rosas et al., 2020***; ***Luppi et al., 2020***), and is a worthwhile avenue for future research.

### Model description and entropy rate estimation

Consider data generated by a stationary, real-valued, discrete-time *d*-dimensional stochastic process 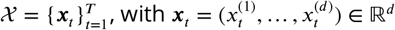, and *T* ∈ ℕ. The core principle of state-space (SS) modelling is to assume that 𝒳 can be modelled as noisy observations of an *m*-dimensional hidden process 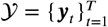, with ***y***_*t*_ ∈ ℝ^m^. Hence, the model is determined by two ingredients: the dynamics of the hidden process 𝒴, and the measurement function that relates 𝒴 with 𝒳. A simple and effective family of models are those with linear dynamics and normally distributed error terms,

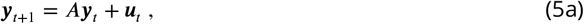

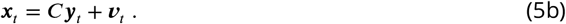

where *A* ∈ ℝ^*m×m*^ is the state transition matrix, *C* ∈ ℝ ^*d*×*m*^ is the observation matrix, and *u*_*t*_, *v*_*t*_ are zeromean white noise processes with covariance matrices 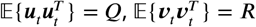, and 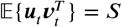.

We calculate the entropy rate of 𝒳 by representing the above SS model in *innovations form*. For this, we consider *“*innovations*”* of the form 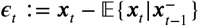, and define 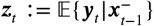, where 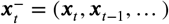 is a shorthand notation for the past trajectory of 𝒳. Then, one can show that the innovations are noise-like, zero-mean i.i.d. variables with covariance 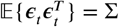.Then, the above SS model can be re-written as

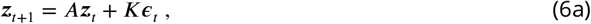

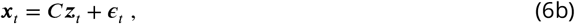

where *K* is typically referred to as the Kalman gain matrix. This is a standard result in time series analysis —for a proof see e.g. ***Durbin and Koopman*** (***2012***, Sec 4.3). With these relationships, one can show that

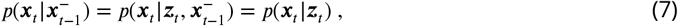

where the first equality follows from the fact that (by definition) 𝓏_*t*_ is a deterministic function of 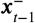, and the second equality from the fact that 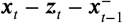 forms a Markov chain. This equation shows one of the key properties of (innovations form) state-space models: that the hidden state 𝓏_*t*_ encapsulates all the knowledge about the signal ***x***_*t*_ one can acquire up to time *t* − 1. Using this, one can plug Eq. (7) into the expression of entropy rate in Eq. (4), and find that

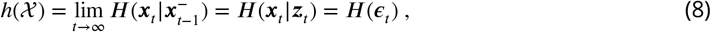

As a final step, given that the innovations are normally distributed, one can compute the entropy rate of 𝒳 using the standard formula for Gaussian distributions (***Cover and Thomas, 2006***, Sec. 8.4):

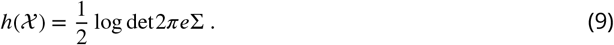

Importantly, before fitting the model and computing CSER we normalise the signals to unit variance. This makes the results invariant to measurement units, prevents biases stemming from differences in total power between two conditions, and allows us to interpret CSER as a function of the proportion of signal variance that is not explained by the past of the signal itself.

It is also worth noting CSER can take negative values. This is because, unlike LZ, CSER estimates the entropy rate of a real-valued distribution (technically called a *differential* entropy rate; ***Cover and Thomas, 2006***). Although differential entropy rates can be negative, their interpretation is the same as their discrete counterpart: higher entropy implies more randomness and less predictability.

In summary, state-space models reduce the daunting task of estimating the entropy rate of the observed process to the estimation the innovations covariance Σ. As we show below, this can be done efficiently with off-the-shelf software, resulting in a flexible, multi-purpose complexity estimator.

### Model selection, convergence speed, and robustness

Given its focus on state-space models, it is no surprise that the core of the CSER computation lies on estimating the SS parameters themselves. To this end, we use the *state-space subspace* (SS-SS) algorithm by ***Van Overschee and De Moor*** (***2012***). There are two parameters we need to specify to run the SS-SS algorithm: the past and future regression horizons (*p, f*), and the model order *m*. Here we present our method for estimating the horizon and model order parameters, without describing the SS-SS algorithm in detail —see ***Van Overschee and De Moor*** (***2012***) for further details.

First, following ***Bauer*** (***2001***), we set the regression horizon using the estimated model order *q* of an AR model, which we fit with the LWR algorithm (***Morf et al., 1978***). We then use the Hannan-Quinn information criterion (***Hannan and Quinn, 1979***) to select the optimal AR order *q*_HQC_, and set *P* = *f* = 2*q*_HQC_. Finally, using these horizons we use the singular value criterion by ***Bauer*** (***2001***) to obtain the optimal state-space model order *m*. This protocol can be implemented with the MVGC toolbox (***Barnett and Seth, 2014***), and results in a method that is flexible, automated, and applicable to various types of data.

To test this procedure, we generated synthetic data from a set of univariate auto-regressive models of order *q* with varying residual variance, and compared the ground-truth entropy rate values with the CSER estimates. We simulated time series of varying length from each model, and computed CSER with the procedure outlined above (Fig. 5, right). For comparison, we did a similar analysis with LZ: we generated synthetic binary data of Markov order *q* (i.e. where *x*_*t*_ is a random boolean function of *x*_*t*−1_, …, *x*_*t*−*q*_ with added noise), and estimated their entropy rate with LZ (Fig. 5, left).

The results show CSER’s clear advantages over LZ as an entropy rate estimator: it is unbiased even for relatively short time series, and variances vanish for longer time series. Even for highly autocorrelated data the variance in CSER drops for time series longer than 10^3^ samples —well within reach of common M/EEG datasets. LZ, in contrast, takes orders of magnitude more samples to converge, especially for time series with long memory (which is the case in most neural data modalities).

On a separate front, one natural question that arises is whether a misestimation of model order *m* may lead to an erroneous estimation of entropy. To address this, we perform a similar test: first, we write an SS model of known order 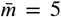 with fixed *A, C, K* parameters; then, we randomly sample a residual covariance matrix from an exponential distribution and simulate the resulting SS model; and finally, we estimate its entropy rate with models of lower order 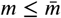 (Fig. 6).

These results again highlight the robustness of the CSER estimator: even when the model order is severely underestimated, CSER is still able to recover the entropy rate of the underlying generative process with high accuracy. Furthermore, CSER is also able to recover the original power spectrum, even if the incorrect model order is used —although it should be noted that if the model order is significantly underestimated the spectrum estimation may suffer.

**Figure 6.**
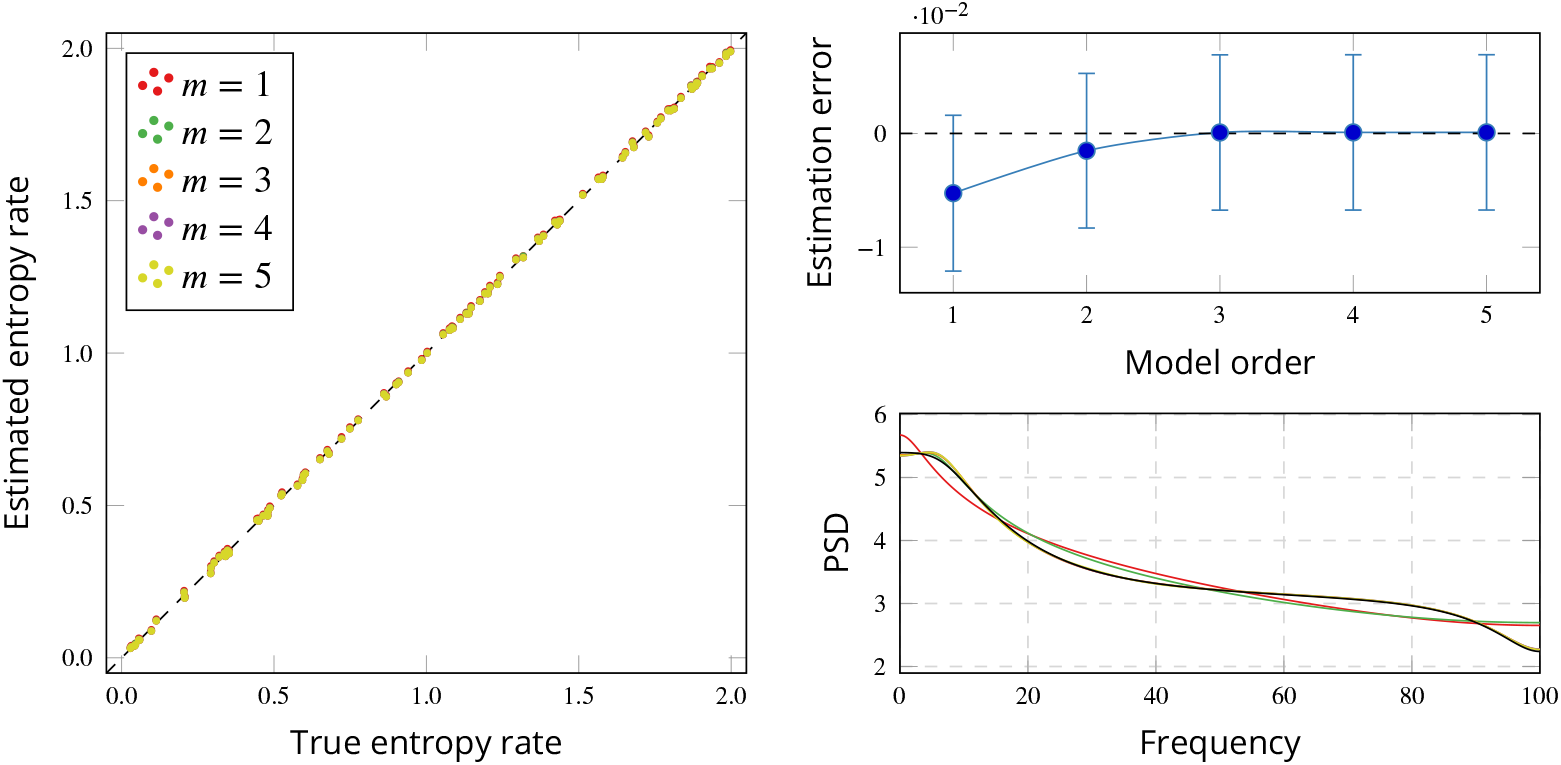
State-space models accurately estimate entropy rate and power spectrum, and are robust to model order selection. Synthetic data was generated from a known state-space model of order 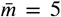, and CSER was computed using different model orders 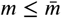. (*left*) True and estimated entropy rate for various model orders. (*top right*) Average estimation error across all runs. (*bottom right*) True (black) and estimated (colours same as left panel) power spectral density (PSD). Although slightly more sensitive (spectrum estimation is visibly inaccurate for *m* ≤ 2), the estimator is still able to recover the true power spectrum with a mis-specified model order.

### Spectral decomposition of CSER

The core element of our spectral entropy rate decomposition is the following expression relating the residual variance and the spectral density of a stationary process, which we state here without proof (***Hannan and Deistler, 2012***, Th. 1.3.2):

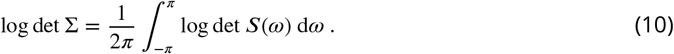

Given the simple expression of the entropy rate in Eq. (9), it is straightforward to re-write the above equation into a spectral decomposition of entropy rate (***Chicharro, 2011***). We do this by adding *d* log(2*πe*) to both sides and using the fact that ***x***_*t*_ is real-valued (and thus (*ω*) = (−*ω*)):

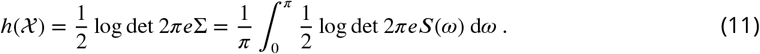

With this expression at hand, we only need to compute the spectral density *S*(*ω*) of the process from the parameters of the state-space model. This is a standard result (***Hannan and Deistler, 2012***), although we rehearse it here for completeness.

We begin by defining *L* as the backshift operator, such that *Lz*_*t*_ = *z*_*t*−1_. With it, we can rewrite Eq. (6a) as *z*_*t*_ = *Az*_*t*−1_ + *K****c***_*t*−1_ = *AL* _*t*_ + *KL****c***_*t*_. Solving for *z*_*t*_ and substituting in Eq. (6b) we obtain the expression of the transfer function of the process, here denoted^12^ by *M*(*L*), as

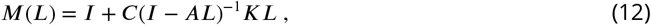

which allows us to write the MA representation of ***x***_*t*_ as a convolution over a white noise process, ***x***_*t*_ = *M*(*L*) * ***ϵ***_*t*_. Now we can simply use the fact that the Fourier transform of a convolution is the product of the Fourier transforms, and arrive at the expression of the spectral density as a function of the transfer function and the residuals’ variance:

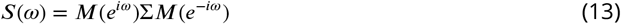

Putting everything together, to compute the results in Fig. 3 we compute the spectral density using Eq. (13) by evaluating Eq. (12) at *e*^*i ω*^, and finally integrate Eq. (11) numerically using the desired (normalised) frequency band as limits for the definite integral.

### Temporally resolved entropy rate

We model the response to the auditory stimulus as a perturbation to an otherwise stationary state-space model. In a nutshell, our method consists of *i*) train a single state-space model using the pre-stimulus baseline of all trials; *ii*) use the trained model to compute prediction errors post-stimulus; and *iii*) compute the log-likelihood of the prediction errors. In the following we describe this procedure in detail. See the Appendix for a technical discussion and validation of this modelling choice.

Recall the ECoG data used here has 720 standard and 720 deviant trials. To train the model, we extract the pre-stimulus baseline (−400 to 0 ms w.r.t. stimulus presentation), stack all 1440 trials together, and train a single state-space model using the model selection procedure described above. This yields a set of (*A, C, K*, I-) parameters, which we leave fixed for the rest of the analysis. Then, for every trial we compute the residuals by performing one-step-ahead predictions —i.e. running the Kalman filter algorithm with fixed parameters (***Durbin and Koopman, 2012***, Sec. 4.3). We compute the residuals for the whole duration of the trial (−400 to 400 ms w.r.t. stimulus presentation), resulting in empirical prediction errors 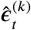, where *t* denotes the time index and *k* the trial number.

Next, we average the prediction errors across trials as 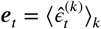, where the average is taken over the standard and the deviant trials separately, resulting in two time series 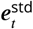 and 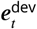 respectively. Finally, using the fact that 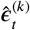 are i.i.d., we know that ***e*** _*t*_ ∼ 𝒩 (0, I-/ *R*), where *R* is the number of trials averaged, so we can compute their log-likelihood as *h*_*t*_ = log 𝒩; (***e***_*t*|_ 0, I-/*R*). The difference 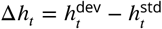 corresponds to the blue curve in Fig. 4.

To determine the statistical significance of the results, we perform a nonparametric cluster test following ***Maris and Oostenveld*** (***2007***). To do this we obtain surrogate samples of 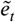 by randomly shuffing the trial labels. We follow the same process to compute surrogate 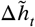 values, against which we compare the observed Δ*h*_*t*_ values to obtain a statistic suitable for the cluster test.

## Acknowledgements

PM and DB were funded by the Wellcome Trust (grant no. 210920/Z/18/Z). FR was supported by the Fellowship Programme of the Institute of Cultural and Creative Industries of the University of Kent. AIL is funded by the Gates Cambridge Trust (OPP 1144). AKS and LB are supported by European Research Council Grant ERC-AdG-CONSCIOUS, project number 101019254, to AKS.

## Appendix Supplementary tables

In this appendix we report the full statistical analysis (mean, standard deviation, Cohen’s *d, t*-score and -value) of subject-level differences across states of consciousness. Tables 1 and 2 correspond to the analyses shown in Figures 2 and 3, respectively.

**Table 1.**
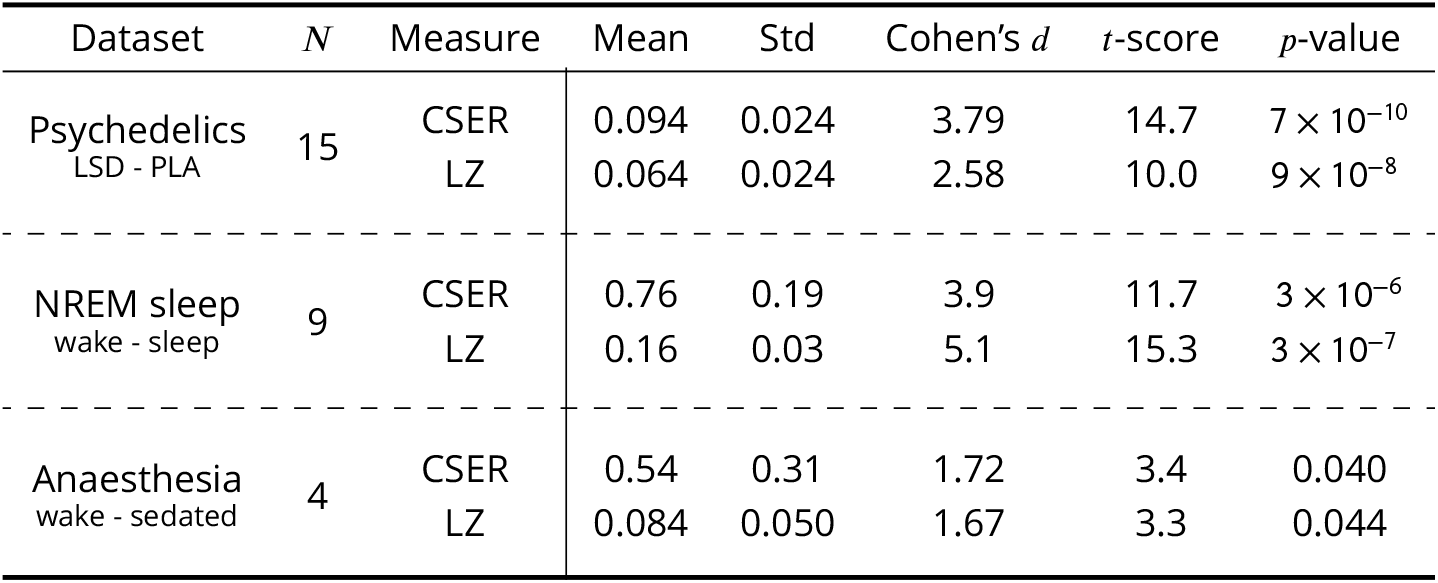
Subject-level differences in broadband CSER and LZ across states of consciousness.

**Table 2.** Subject-level differences in spectral CSER across states of consciousness. Frequency bands used are *δ*: 1–4 Hz, *θ*: 4–8 Hz, *α*: 8–14 Hz, *β*: 14–25 Hz, *γ*: >25 Hz.

| Dataset | *N* | Band | Mean | Std | Cohen's *d* | *t*-score | *p*-value |
| --- | --- | --- | --- | --- | --- | --- | --- |
| Psychedelics<br>LSD - PLA | 15 | $\delta$ | -0.0025 | 0.0022 | -1.14 | -4.4 | $5 \times 10^{-4}$ |
| | | $\theta$ | -0.0047 | 0.0024 | -1.88 | -7.3 | $3 \times 10^{-6}$ |
| | | $\alpha$ | -0.0083 | 0.0038 | -2.14 | -8.3 | $8 \times 10^{-7}$ |
| | | $\beta$ | -0.0058 | 0.0067 | -0.85 | -3.3 | 0.004 |
| | | $\gamma$ | 0.11 | 0.03 | 3.28 | 12.7 | $4 \times 10^{-9}$ |
| NREM sleep<br>wake - sleep | 9 | $\delta$ | -0.017 | 0.005 | -3.21 | -9.6 | $1 \times 10^{-5}$ |
| | | $\theta$ | -0.0068 | 0.008 | -0.83 | -2.5 | 0.03 |
| | | $\alpha$ | 0.037 | 0.009 | 4.00 | 12.1 | $2 \times 10^{-6}$ |
| | | $\beta$ | 0.080 | 0.01 | 7.37 | 22.1 | $2 \times 10^{-8}$ |
| | | $\gamma$ | 0.67 | 0.17 | 3.80 | 11.4 | $3 \times 10^{-6}$ |
| Anaesthesia<br>wake - sedated | 4 | $\delta$ | -0.003 | 0.005 | 0.50 | -1.0 | 0.39 |
| | | $\theta$ | 0.022 | 0.007 | 3.15 | 6.3 | 0.008 |
| | | $\alpha$ | 0.06 | 0.012 | 4.45 | 8.9 | 0.003 |
| | | $\beta$ | 0.11 | 0.031 | 3.54 | 7.1 | 0.006 |
| | | $\gamma$ | 0.37 | 0.27 | 1.32 | 2.6 | 0.07 |

**Table 3.** Subject-level differences in spectral CSER across states of consciousness, controlled for gamma coherence. Results correspond to the regression coefficients for conscious state in the linear model ‘CSER State + Coherence’. Coherence was calculated using the Fieldtrip toolbox (***Oostenveld et al., 2011***) with a multitaper Fourier transform method, and averaged across all pairs of channels. Effect size *d* here was calculated as suggested by ***Feingold*** (***2013***), taking the regression coefficient divided by the residual standard deviation of the full model. Frequency bands used are *δ*: 1–4 Hz, *θ*: 4–8 Hz, *α*: 8–14 Hz, *β*: 14–25 Hz, *γ*: >25 Hz.

| Dataset | $N$ | Band | Mean | Std | Cohen’s $d$ | $t$-score | $p$-value |
| --- | --- | --- | --- | --- | --- | --- | --- |
| Psychedelics<br>LSD vs PLA | 15 | All | 0.094 | 0.012 | 3.83 | 7.47 | $5 \times 10^{-8}$ |
| | | $\delta$ | -0.0025 | 0.0010 | -0.96 | -2.56 | 0.016 |
| | | $\theta$ | -0.0047 | 0.0010 | -1.71 | -4.53 | $1 \times 10^{-4}$ |
| | | $\alpha$ | -0.0083 | 0.0014 | -2.21 | -5.8 | $3 \times 10^{-6}$ |
| | | $\beta$ | -0.0058 | 0.0031 | -0.69 | -1.83 | 0.07 |
| | | $\gamma$ | 0.12 | 0.016 | 2.78 | 7.35 | $6 \times 10^{-8}$ |
| NREM sleep<br>wake vs sleep | 9 | All | 0.69 | 0.07 | 5.17 | 9.76 | $6 \times 10^{-8}$ |
| | | $\delta$ | -0.019 | 0.001 | -6.15 | -11.6 | $6 \times 10^{-9}$ |
| | | $\theta$ | -0.0075 | 0.003 | -1.38 | -2.6 | 0.02 |
| | | $\alpha$ | 0.039 | 0.003 | 6.37 | 12.0 | $4 \times 10^{-9}$ |
| | | $\beta$ | 0.082 | 0.006 | 6.48 | 12.2 | $3 \times 10^{-9}$ |
| | | $\gamma$ | 0.60 | 0.07 | 4.57 | 8.6 | $3 \times 10^{-7}$ |
| Anaesthesia<br>wake vs sedated | 4 | All | 0.54 | 0.21 | 2.17 | 2.59 | 0.048 |
| | | $\delta$ | -0.003 | 0.002 | -1.15 | -1.37 | 0.23 |
| | | $\theta$ | 0.022 | 0.004 | 4.24 | 5.1 | 0.003 |
| | | $\alpha$ | 0.056 | 0.01 | 4.63 | 5.5 | 0.002 |
| | | $\beta$ | 0.11 | 0.02 | 3.58 | 4.3 | 0.008 |
| | | $\gamma$ | 0.37 | 0.17 | 1.75 | 2.1 | 0.09 |

## Appendix LZ is a measure of entropy, not algorithmic complexity

A significant portion of the neuroscience literature that employs LZ argues that it is somewhat akin to the Kolmogorov-Chaitin algorithmic complexity (***Sitt et al., 2014***; ***Casali et al., 2013***). This is by no means an unmotivated belief, as the original ideas of Lempel and Ziv were certainly motivated by Kolmogorov’s work (***Lempel and Ziv, 1976***), and well-known references promote the use of LZ as an upper bound to Kolmogorov-Chaitin algorithmic complexity (***Li et al., 2008***). Although this is not technically incorrect, in what follows we argue that this can lead to undesirable misinterpretations.

Recent investigations have shown that, despite superficial similarities, Shannon’s entropy and algorithmic complexity are fully dissociated (***Zenil et al., 2019***). In particular, while Shannon’s entropy can always be used as upper bound of the algorithmic complexity, this bound can be infinitely inaccurate.^13^ Furthermore, there is reason to be skeptical of alleged estimators of algorithmic complexity from experimental data. Kolmogorov’s *Invariance* (or *Universality*) *Theorem* states that the Kolmogorov complexity of a sequence ***x*** read with two different Turing machines differs by a constant independent of ***x*** (***Cover and Thomas, 2006***, Ch. 14). An important corollary for practical applications is that said constant can be *arbitrarily large* for any finite sequence. For practical purposes, that means that a measured difference between two conditions may be a genuine difference in algorithmic complexity, or may just be a Turing machine mismatch —which has a substantially different interpretation. This is not to say that algorithmic complexity cannot be estimated from data —see e.g. ***Zenil et al. (2018***) –, although there is great nuance involved which cannot be simply swept under the rug of the LZ algorithm.

In addition to the concerns above, there is one more hurdle to the Kolmogorov complexity interpretation of LZ: it relies on the fundamental assumption that the sequence must be exactly reproduced by a Turing machine (***Cover and Thomas, 2006***). This is a very strong assumption to make regarding neural dynamics, which clashes with contemporary accounts of the brain as a nonlinear stochastic system (***Deco et al., 2009***). For these reasons, we believe the interpretation of LZ as entropy rate is both more mathematically principled and more parsimonious, and thus should be preferred when interpreting empirical results.

## Appendix Alternatives for entropy estimation in non-stationary data

In general, modelling of ERPs is a difficult task due to the strongly non-stationary nature of event-related data (***Ding et al., 2000***). As mentioned in the Methods section, the time-resolved entropy analysis shown in Fig. 4 was performed assuming a constant model subject to a non-stationary perturbation. For completeness, here we discuss an alternative approach: one could leverage the trial structure of the data by fitting SS models using all trials and sliding temporal windows, and computing CSER for each of them.

In essence, the approach used in the main text corresponds to modelling the ERP as nonstationary innovations in a stationary model, and the approach introduced here corresponds to stationary innovations in a non-stationary model. In principle, these two alternatives can be pitted against each other through standard model selection tests, e.g. via a likelihood ratio test. However, the likelihood function of a non-stationary state-space model is complicated, and therefore this approach is difficult in practice.

Nonetheless, as an example of this approach and as a validation of the approach from the main text, we performed a sliding-window CSER analysis on the same oddball task data as before. For this, we processed the data following recommendations by ***Ding et al. (2000***) (including time-wise ensemble demeaning and normalisation) and computed CSER in sliding windows of 20 samples. We performed this analysis both with and without filtering and downsampling, to obtain a more granular picture of the CSER time-course (Fig. 7).

**Figure 7.**
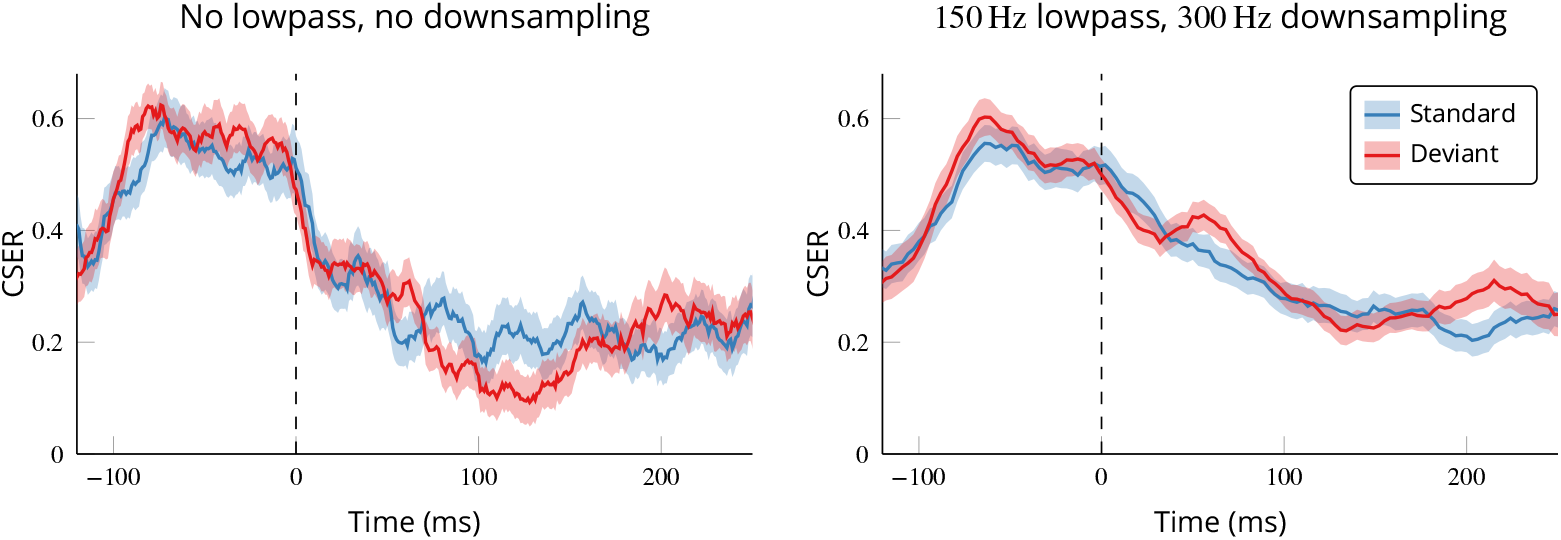
Sliding-window CSER estimates show no signi1cant differences between standard and deviant tones. Results are shown before (*left*) and after (*right*) applying a 150 Hz lowpass filter to the data and downsampling it to 300 Hz. A standard cluster test found no significant differences between conditions in either case.

The most visible result from this analysis is that indeed there is temporal structure in the sliding-window CSER across the trial, most noticeably a sharp dip around the stimulus time. This could be interpreted as the stimulus dominating the signal and suppressing endogenous noise; or, in more cognitive terms, as there being some uncertainty about the precise stimulus timing which is resolved as soon as the stimulus is presented.

Strictly speaking, this means the assumptions behind the model from Fig. 4 are not fully correct, since at least one parameter (Σ) changes over the course of the trial. However, when comparing the standard and deviant CSER values, there are no significant differences between the two types of stimuli surviving a cluster test (***Maris and Oostenveld, 2007***). Therefore, although it does seem likely that model parameters change throughout the course of the trial, this change does not account for the difference between stimuli found in Fig. 4.

To summarise, we have described two different ways in which a time-resolved entropy rate time course can be obtained from multi-trial data leveraging state-space models. These make different assumptions, and although they are difficult to test rigorously, they may have different advantages in different situations.

## Footnotes

1 Technically, this applies only under certain conditions. The process must be ergodic (and strict-sense stationary), so that its joint probability distribution does not change over time —i.e. for any set of indices { _1_, …,} the process satisfies that 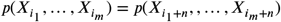 for all Although strict, this is a common requirement in time series analysis methods.

2 Note that this normalisation applies only to the LZ76 algorithm proposed in ***Lempel and Ziv*** (***1976***). Later versions (e.g. LZ77 or LZ78) also converge to the entropy rate but need other normalisation values.

3 Although see contradictory reports in ***Pal et al. (2020***).

4 Note that this is different from simply pre-filtering the signals at specific frequency bands and then computing LZ (or CSER) on the filtered sequence, which has no relation to the broadband LZ (or CSER).

5 Though they later relaxed their claims on the topic.

6 For example, in the case of ***Timmermann et al. (2019***), the LZ changes are driven by the pharmacokinetics of DMT that take place at a much slower time scale than neural activity.

7 As an analogy, consider the case of an orbiting planet being hit by an asteroid. The planet’s trajectory is highly predictable (knowing Kepler’s laws), up to the moment of impact —when a high prediction error would occur in a hypothetical observer. After the impact, the ability to predict is restored, as Kepler’s laws continue to apply. Importantly, the prediction error (increase in instantaneous entropy rate) *precedes* the time when the new orbit maximally deviates from the original one (akin to difference in activity).

8 More technically, we say that state-space models are *closed* with respect to several transformations that neural data typically goes through. For example, if a system of variables following a state-space model is linearly mixed (e.g. with a forward or inverse model for source reconstruction), or is temporally or spatially subsampled, the resulting system is also a state-space model. The same is not true of other time series models, such as auto-regressive models.

9 Technically, this is because the HRF may not be a *‘*minimum-phase’ filter —although this claim has been contested (***Barnett and Seth, 2015***). A separate concern is that fMRI operates at timescales much slower than the underlying neural process, which makes drawing conclusions about neural dynamics from fMRI a difficult task (***Barnett and Seth, 2017***).

10 This is a direct consequence of the Asymptotic Equipartition Property, and most fundamentally from the Shannon-McMillan-Breiman theorem for stationary ergodic processes (***Cover and Thomas, 2006***, Ch. 3).

11 *‘*Intrinsic’ here means that it is a property of the stochastic process itself, and does not depend on which model one may choose to predict the next value of the signal (since it is the minimum error achievable by any model).

12 Unfortunately, the standard symbol for both entropy and the transfer function is the letter *H*. Although it should be clear from the context, to avoid confusion we adopt the non-standard symbol *M* for the transfer function.

13 As a simple example of this, consider strings generated via this short snippet in the Python programming language: `negate = lambda x: 1 - x; reduce(lambda s,r: s + list(map(negate, s)), range(T), [0])` The usual dictionary-based implementation of LZ yields large complexity values, which diverge to infinity as T grows.

